# Auxin-dependent acceleration of cell division rates regulates root growth at elevated temperature

**DOI:** 10.1101/2022.06.22.497127

**Authors:** Haiyue Ai, Julia Bellstaedt, Kai Steffen Bartusch, Lennart Eschen-Lippold, Steve Babben, Gerd Ulrich Balcke, Alain Tissier, Bettina Hause, Tonni Grube Andersen, Carolin Delker, Marcel Quint

## Abstract

Roots are highly plastic organs enabling plants to acclimate to a changing below-ground environment. In addition to abiotic factors like nutrients or mechanical resistance, plant roots also respond to temperature variation. Below the heat stress threshold, *Arabidopsis thaliana* seedlings react to elevated temperature by promoting primary root growth, possibly to reach deeper soil regions with potentially better water saturation. While above-ground thermomorphogenesis is enabled by thermo-sensitive cell elongation, it was unknown how temperature modulates root growth. We here show that roots are able to sense and respond to elevated temperature independent of shoot-derived signals. A yet unknown root thermosensor seems to employ auxin as a messenger to promote primary root growth. Growth is primarily achieved by accelerating cell division rates in the root apical meristem, likely maintained via temperature-sensitive organization of the polar auxin transport system. Hence, the primary cellular target of elevated ambient temperature differs fundamentally between root and shoot tissues, while the messenger auxin that relays temperature information to elongating or dividing cells, respectively, remains the same.

## INTRODUCTION

With increasing concern over global warming, awareness of its impact on living organisms has steadily grown in the past 30 years. Plants can acclimatize to warm temperatures with wide-ranging physiological and morphological adjustments. These acclimation processes can enhance plant fitness, but high ambient temperatures still exert a negative effect on plant growth and development, and likewise on crop yields (Lippmann *et al*., 2019). Thermomorphogenesis describes these acclimations and thereby the effect of ambient temperature on plant morphogenesis (Delker *et al*., 2014). In the model plant *Arabidopsis thaliana*, hypocotyl elongation and increased leaf hyponasty are among the earliest thermomorphogenic changes in response to elevated temperatures (Quint *et al*., 2016; Balasubramanian and Casal, 2019). Such morphological adjustments result in an open rosette structure with increased ventilation enabling evaporative cooling which effectively reduces leaf temperatures (Crawford *et al*., 2012). Across land plants, the ecophysiological advantage of these acclimation responses may be to confer some sort of metabolic plasticity, mitigating the effects of high temperature (Ludwig *et al*., 2021). The major signaling hub controling these morphological responses in *A. thaliana* is the basic helix-loop-helix transcription factor PHYTOCHROME-INTERACTING FACTOR 4 (PIF4) (Koini *et al*., 2009). In concert with downstream hormonal action (reviewed in Castroverde and Dina, 2021), PIF4 transcriptionally regulates genes promoting cell elongation. PIF4 itself is regulated by various thermosensors. At low temperature, PIF4 is repressed by numerous mechanisms that affect PIF4 expression, protein levels and activity (reviewed in Delker et. al. 2022). Important thermosensors that directly repress PIF4 are PHYTOCHROME B (phyB, Jung *et al*., 2016; Legris *et al*., 2016) and EARLY FLOWERING 3 (ELF3, Jung *et al*., 2020). High temperature relieves this inhibition and induces *PIF4* transcription and other PIF4 regulatory mechanisms (reviewed in Delker et al. 2022), opening the route to thermo-induced elongation growth. So far, these discoveries describe thermomorphogenic responses only in shoot tissues.

Similar elongation responses occur in roots (e.g., Quint *et al*., 2005; Quint *et al*., 2009; Hanzawa *et al*., 2013; Wang *et al*., 2016; Ibanez *et al*., 2017; Martins *et al*., 2017; Feraru *et al*., PNAS, 2019; Gaillochet *et al*., 2020; reviewed in de Lima *et al*., 2021). In contrast to the shoot, the ecophysiological basis of these responses is less clear for underground tissues. Possibly, thermo-induced root growth is coupled to drought responses, enabling the plant to reach deeper soil layers with presumably better water saturation (Martins *et al*., 2017; Ludwig *et al*., 2021). However, our mechanistic understanding of temperature-sensitive root growth is rather fragmentary. Given that roots grow below the surface, it seems unlikely that temperature-sensitive parts of photomorphogenesis pathways, which are central to shoot thermomorphogenesis (reviewed in Qi et al., 2022), represent the primary temperature sensing pathway also in roots. Several phytohormones have been suggested to play a role in root thermomorphogenesis. While the dependence of temperature-induced root elongation on auxin metabolism, transport and signaling is established (reviewed in de Lima *et al*., 2021), proposed roles for brassinosteroids (Martins *et al*., 2017), ethylene and gibberellic acid (Fei *et al*., 2017, 2019) would benefit from additional mechanistic insight. A general challenge for assigning specific roles of phytohormones to physiological processes is often the pleiotropic nature of many hormonal mutants. Such mutants tend to display severe root growth defects already at control temperatures, potentially masking temperature-specific effects. Therefore, conditional mutants would be most informative.

In this study, we first ask whether roots are autonomously able to sense and respond to elevated temperatures or whether they require shoot-root long distance communication as proposed previously (Gaillochet *et al*., 2020). We then aim to uncover the molecular mechanism that is responsible for temperature-induced root elongation. We show that, while the messenger delivering temperature information to the cells seems to be the same, the cellular process enabling thermo-responsive root growth differs fundamentally from the one regulating thermomorphogenesis in above-ground tissues.

## RESULTS

### Root growth dynamics in response to different temperatures

Based on recently published work from other labs (e.g., Martins *et al*., 2017; Feraru *et al*., 2019; Gaillochet *et al*., 2020; Lee *et al*., 2021) and our own previous comprehensive phenotypic analysis (including root growth) of various *A. thaliana* accessions grown in a range of different ambient temperatures (Ibanez *et al*., 2017), we tried to better understand how temperature specifically affects root growth. In the temperature range investigated (20°C *vs*. 28°C), elevated ambient temperature obviously promoted primary root elongation in *A. thaliana* accession Col-0 (Fig. 1A-C), but also across species (Fig. 1D-E), demonstrating that temperature-induced root elongation is a universal response. Daily measurements of root length in *A. thaliana* suggested that growth differences were absent in the first four days of cultivation (Fig. 1B). To substantiate this observation, we captured root growth rates continuously and in real-time between days 2 to 7 after germination. Infrared imaging enabled hourly documentation of growth behavior also in darkness. Similar to the end point analysis presented in Fig. 1B, we observed differential growth rates only after day 4 (Fig. 1C). Independent of temperature, root growth rates displayed a pronounced diurnal fluctuation (Fig. 1C). Under the long day photoperiods applied, root growth rates increased in late afternoon and peaked at the end of the night, which was mirrored by high temperature hypocotyl growth patterns in the same plants (Fig. EV1A). These data indicate that (under our conditions) root growth rates increase during early seedling development, but temperature sensitivity of growth rates seems to be gated during the first few days.

**Figure 1.**
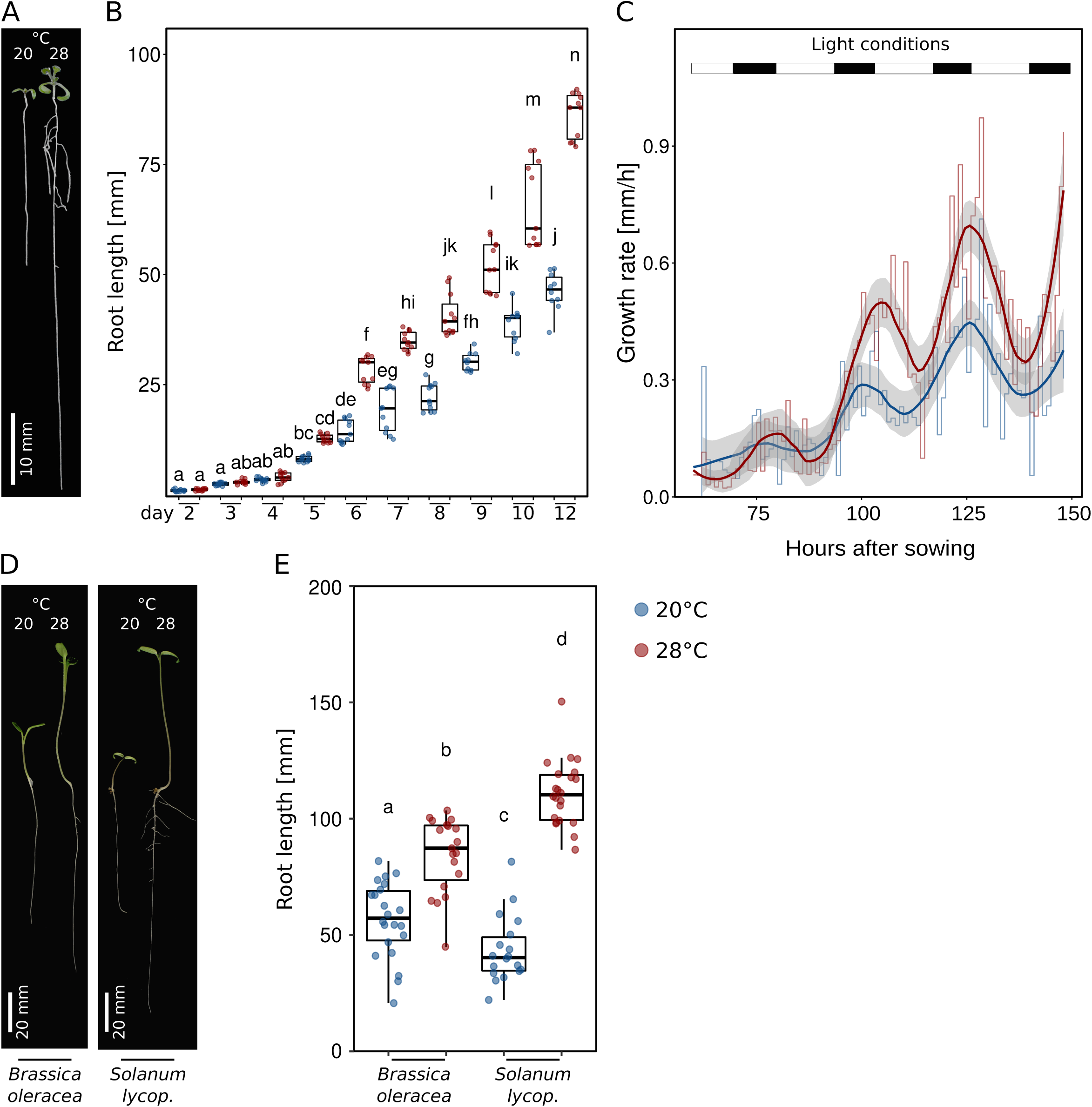
Root growth dynamics in response to different temperatures. **A** Arabidopsis seedlings grown for 7 days in LD at 20°C or 28°C. **B** Root lengths of seedlings from day 2 to day 12 after sowing (grown as in A). **C** Growth rates of seedlings between days 2-7 were assessed by hourly infrared real-time imaging. Mean growth rates are shown as step-wise lines (n = 8) that were fitted with a ‘loess’ smoothing function shown as solid lines and the respective 95 % confidence intervals as grey ribbons. Long-day lighting is schematically depicted by white (light) and black (dark) bars on the top. **D** Seedlings of *Brassica oleracea* and *Solanum lycopersicum* (*lycop*.) grown for 7 days at 20°C or 28°C. **E** Root lengths of 7 days-old seedlings grown at 20°C or 28°C (n = 18-22). B,E Boxplots show medians, interquartile ranges and min-max values with individual data points superimposed as colored dots. Different letters denote statistical differences at P < 0.05 assessed by 2-way ANOVA and Tukey’s HSD *posthoc* test.

### Roots are independently sensing and responding to differences in ambient temperature

We had previously suggested that roots are able to autonomously sense and respond to temperature without the need for shoot-derived signals (Bellstaedt *et al*., 2019). However, Gaillochet *et al*. (2020) proposed the necessity of shoot-to-root communication for root thermomorphogenesis. To revisit this question and to assess whether or not the root response requires such long distance signals from a temperature-sensing shoot, we first tested temperature-induced root elongation assays with excised roots (only the root, no shoot attached) in Col-0. In confirmation of our previously published data (Bellstaedt *et al*., 2019), we found that detached roots were not only able to continue growing in the absence of shoot tissue, they even elongated more at 28°C than at 20°C (Fig. 2A). This suggests that roots can be regarded as autonomous in terms of sensing and responding to temperature cues. This behavior was similar in *Brassica oleracea* and *Solanum lycopersicum* (Fig. EV1B), indicating its conservation across species.

**Figure 2.**
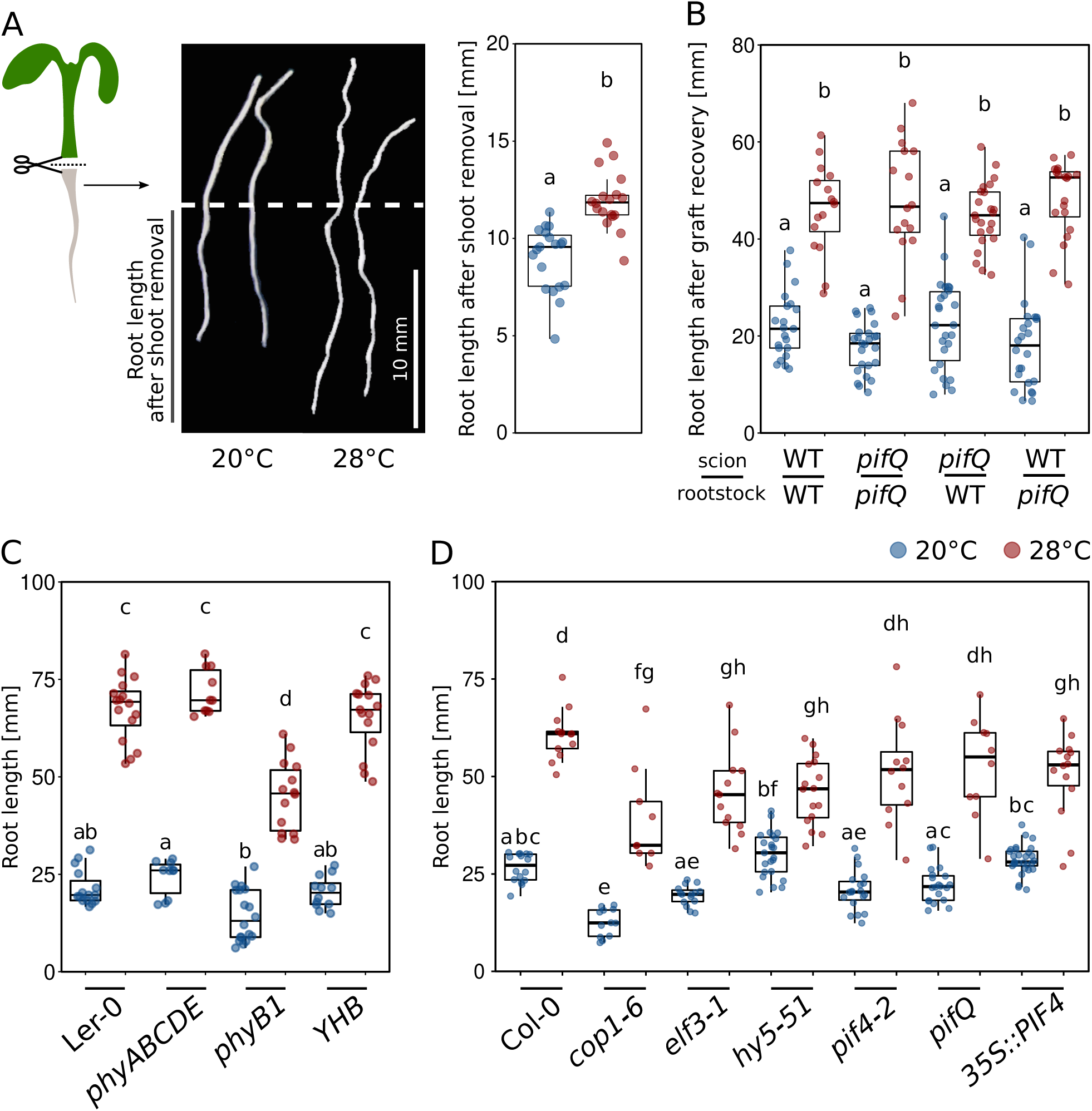
The root is able to autonomously sense temperature and respond to it. **A** Elongation responses of detached roots. Shoots were removed from 4 days-old seedlings grown at 20°C and detached roots were grown for additional 4 days at 20°C or 28°C (n = 18-19), scale bar = 10 mm. **B** 9 days-old seedlings were hypocotyl-grafted, recovered for 7 days and then cultivated at 20°C or 28°C for additional 7 days (n = 16-26). **C**,**D** Root growth assay of seedlings grown for 7 days (C n= 10-18, D n= 9-29) at the indicated temperatures. **A-D** Boxplots show medians, interquartile ranges and min-max values with individual data points superimposed as colored dots. Different letters denote statistical differences at P < 0.05 as assessed by one-way (A) or two-way (B-D) ANOVA and Tukey’s HSD *posthoc* test.

As this assay is rather ‘rude’, we next performed a likewise invasive but certainly less drastic hypocotyl micrografting assay to substantiate these results, also in different genetic backgrounds (Fig. EV2A). Here, a shoot mutant that is unable to transduce temperature cues would be informative, as in such a genetic background the relay of temperature-induced signals from the shoot to the root is unlikely. For these experiments, we first used the essentially temperature-blind quadruple *pifQ* (*pifQ* = *pif1-1 pif3-7 pif4-2 pif5-3*) mutant. Fig. 2B shows that all possible *pifQ*/wt scion > rootstock combinations showed a wild type-like root elongation in response to elevated temperature. This behavior was, not surprisingly, mirrored by a *pif4-2* single mutant (Fig. EV2B). Likewise, *hy5-51* mutant shoots on wild type rootstocks behaved like wild type, arguing against a role for shoot localized or shoot-derived HY5 in root thermomorphogenesis (Fig. EV2C). Only *hy5-51* shoots grafted onto *hy5-51* rootstocks showed shorter roots at both temperatures, which is in line with the decreased temperature response of intact *hy5-51* seedlings (Fig. EV2C and Lee *et al*., 2021; Gaillochet *et al*., 2020). The only line with a potential shoot-to-root effect in these experiments was the shoot thermosensory mutant *phyB-9*. While wild type shoots on *phyB-9* rootstocks did not differ from the wild type, graft combinations including *phyB-9* shoots displayed significantly shorter roots at high temperature when compared to graft combinations with wild type shoots (Fig. EV2D). However, these grafting combinations were still able to respond to the temperature stimulus, suggesting a rather minor role for shoot-localized or -derived phyB in this process. It has to be noted that the growth conditions and times needed for grafting assays (Fig. EV2A) differ from the experimental setup for intact seedlings. Together with the still invasive character of micrografting and some hypocotyl tissue that remains with the rootstock, this should be taken into account when interpreting these data. Furthermore, we are aware that at this point there is no true negative control available showing a severe root elongation defect in response to temperature. Taken together, although a shoot-derived function of phyB cannot be ruled out, the thermo-responsive behavior of micrografted seedlings with thermosensing/-signaling defective scions on wild type rootstocks largely supports an autonomous character of roots in terms of their capability to sense and respond to elevated ambient temperature.

In intact seedlings we found that none of the shoot thermomorphogenesis mutants tested (*phy* loss- and gain of function lines, *cop1-6, elf3-1, pif* loss of function and overexpression lines) showed a root response defect that was remotely reminiscent of their severe shoot phenotypes (Fig. 2C,D; Delker *et al*., 2014; Jung *et al*., 2016; Legris *et al*., 2016). While we did observe statistically significant differences for selected mutant lines in comparison to the wild type (see also Gaillochet *et al*., 2020), these differences were mostly subtle. All tested lines were obviously still able to sense elevated temperatures and respond with root growth promotion. Again, absence of a temperature unresponsive root mutant complicates interpretation of these data because of the unavailability of such a negative control. It has to be noted, however, that others have provided solid evidence, suggesting that at least HY5 seems to play a role in root thermomorphogenesis, albeit as a positive regulator in contrast to its repressive function in the shoot (Gaillochet *et al*., 2020; Lee *et al*., 2021). Taken together, under the expectation of severe phenotypes in loss-of-function backgrounds of important signaling components, we conclude so far that roots are able to autonomously sense and respond to temperature, most likely via a signaling pathway that is different from the pathway regulating the shoot temperature response. While a role for phytochrome-dependent thermo-signaling, which dominates shoot thermomorphogenesis, cannot be generally ruled out, this role is likely of secondary or indirect nature in thermo-responsive root growth.

### Temperature-induced root growth is largely due to promotion of cell proliferation

Theoretically, temperature-induced primary root elongation may be a consequence of temperature-promoted cell elongation (as in the shoot), temperature-promoted cell division, or a combination thereof. To assess the potential of both processes, we first compared the number of hypocotyl or radicle/root cells in mature embryos *vs*. seven days-old seedlings (Col-0). We observed that the number of cortical hypocotyl cells along a longitudinal cell file from the root-shoot junction to the shoot apical meristem was only marginally larger in seedlings compared to mature embryos, with a significant but minor temperature effect (Fig. EV3A). This incremental increase of number of hypocotyl cells during early seedling development practically leaves only cell elongation as the primary mechanism of vertical organ growth in response to warmth, while cell division can likely be neglected. In roots, the picture changed dramatically. The embryonic radicle consists of only a few cells, while cell division generates hundreds of new root cells post germination along a longitudinal cell file (Fig. EV3B), and therefore thousands in the whole 3-dimensional root. Furthermore, the number of root cells further increased at 28°C. This indicates that cell division contributes substantially to thermo-responsive root growth.

We therefore investigated cellular growth patterns and counted and measured root cells along a longitudinal cell file from the quiescent center to the root-shoot junction in wild type (Col-0) seedlings. Fig. 3A shows temperature dependency of cell proliferation and its consequences for root elongation. It had previously been reported that in high temperatures the meristematic zone slightly decreased in size (Yang *et al*., 2017; Feraru *et al*., 2019). In our assays, we did not observe major effects of temperature on the number and length of cells neither in meristematic nor in elongation zones of roots in five days-old seedlings (Fig. 3B). However, a daily inspection of meristematic and elongation zone length showed that root apical meristems of high temperature grown seedlings were slightly larger during the first few days until they leveled out after six days (under our conditions). Meristems of low temperature grown seedlings continued to increase in length also after six days (Fig. EV4A). In general, the differences in meristem size between temperatures seemed subtle. A similar picture emerged for the elongation zone (Fig. EV4B). In contrast, we did observe a substantially larger differentiation zone (defined as distal to the elongation zone up to the root-shoot junction, Fig. EV4C) due to an increased cell number in this zone (Fig. S4D) in 28°C grown seedlings compared to 20°C grown seedlings. Cell lengths in the same zone remained, however, rather stable (Figs 3A, EV4E). Together, this indicates that in the absence of notable differences in (i) cell length throughout the root, and (ii) cell numbers in meristematic and elongation zones, the only reasonable explanation for temperature-induced root elongation is a higher rate of cell division in the root apical meristem, resulting in more cells being released to the elongation zone in a defined time period.

**Figure 3:**
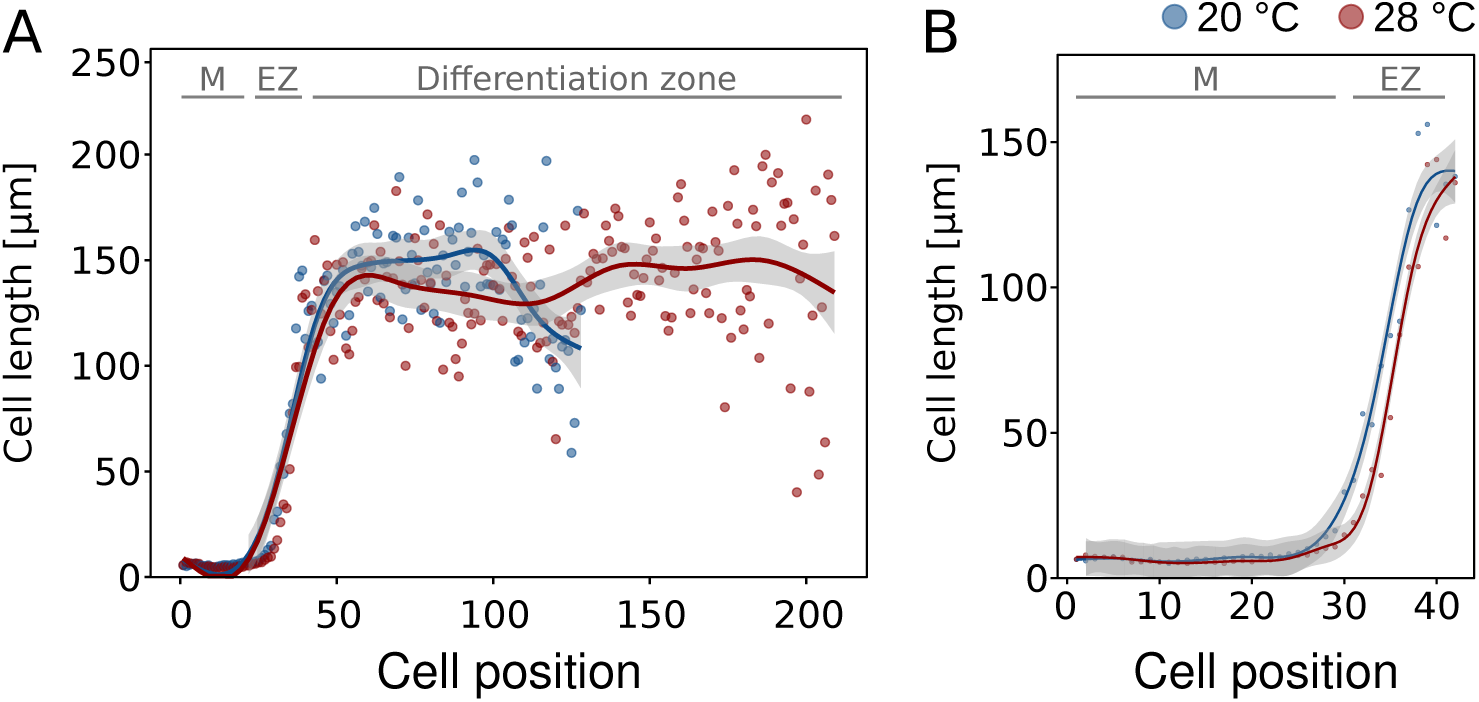
Temperature effects on root cell size and numbers. **A** Cell lengths in consecutive cortex cell files of 5 days-old seedlings starting from the root tip (quiescent center = position 1) spanning the meristem (M), elongation zone (EZ), and differentiation zone up to the root-shoot junction. Individual dots represent mean cell lengths (n = 8), lines show a fitted smoothing function (generalized additive models) with the 95 % confidence intervals shown in light grey ribbons. **B** Close-up view of the first 43 cells comprising M and EZ of the data shown in A.

Hence, while shoot thermomorphogenesis is driven by cell elongation, root thermomorphogenesis seems to be dominated by temperature effects on cell division, representing a fundamental mechanistic difference between the two organs. This suggests that temperature sensing and signaling is regulated via different pathways in roots and shoots, which is supported by largely different transcriptome responses to elevated temperature in root *vs*. shoot tissues we observed previously (Bellstaedt *et al*., 2019).

### Elevated temperature promotes cell division rates in the root apical meristem

To test whether cell division rates increase at elevated ambient temperatures, we quantified the number of meristematic cells that are either entering the cell cycle (S-phase), transitioning from G2 to mitosis or are actively dividing (M-phase). 5-ethynyl-2’-deoxyuridine (EdU), a thymidine analog, is widely used in DNA proliferation assays. EdU is incorporated into newly synthesized DNA and stains meristematic cells in the S-phase. When EdU labeled cells divide, each daughter cell also contains EdU labeled nuclei. Consequently, an increased number of labeled cells indicates a higher cell division rate. Quantifying the pCYCB1::CYCB1-GFP reporter expressed in cytrap lines (Yin *et al*., 2014) enables monitoring G2/M phase transition. And lastly, 4′,6-diamidino-2-phenylindole (DAPI) is a fluorescent stain that binds strongly to adenine-/thymine-rich regions in DNA allowing identification of cells in mitosis (M-phase). The experiments were conducted with five days-old seedlings, which is after the temperature-response gates opened as shown above (Figs 1B,C). Wild type Col-0 seedlings were grown in either constant 20°C or constant 28°C. Fig. 4A shows that high temperature significantly increased the number of cells entering the cell cycle. Likewise, cells in G2/M transition (Fig. 4B) and those actively dividing (Fig. 4C) increased significantly in high temperature. This supports the hypothesis stated above that higher temperature increases cell division rates.

**Figure 4:**
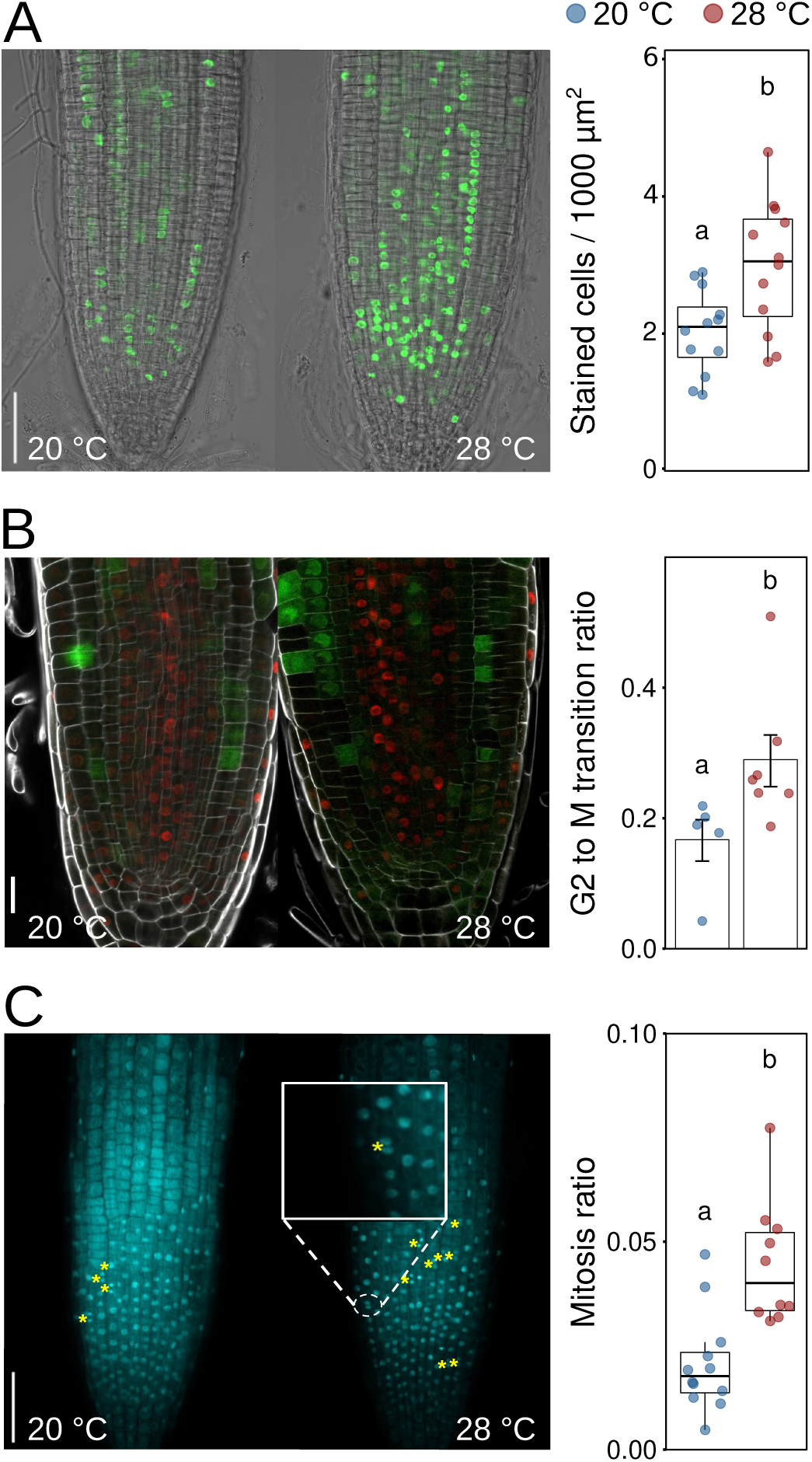
Warm temperature accelerates the cell cycle. Different stages of the cell cycle were assayed in 5 days-old Arabidopsis seedlings grown in LD at 20°C or 28°C. **A** EdU staining marks cells that were in S-phase during the 1h incubation time prior to microscopic imaging.Scale bar = 50 µm. **B** Cytrap lines highlight cells in G2 to M phase transition. Scale bar = 20 µm. **C** DAPI staining was used to identify cells currently in mitosis (examples are highlighted by yellow asterisks. Scale bar = 50 µm. **A-C** Boxplots show medians, interquartile ranges and min-max values with individual data points superimposed as colored dots (n = 5-14 individual roots). Different letters denote statistical differences at P < 0.05 as assessed by one-way ANOVA and Tukey’s HSD *posthoc* test.

### Auxin transfers temperature information to the cell cycle

While these data indicated that the increase of cell division rates is an important driver of root growth at elevated temperatures, it remained unclear how temperature information is perceived in the root and how it reaches the cell cycle. A likely intermediate signal between a yet unknown root thermosensor and the cell cycle that has been shown to be involved in both root thermomorphogenesis (Hanzawa *et al*., 2013; Wang *et al*., 2016; Feraru *et al*., 2019; Gaillochet *et al*., 2020) and cell cycle regulation (reviewed in Perrot-Rechenmann, 2010; del Pozo and Manzano, 2014) is the phytohormone auxin.

We first followed a pharmacological approach to test whether decreasing free indole-3-acetic acid (IAA) levels had an effect on temperature-induced root elongation. Fig. 5A shows that increasing concentrations of the IAA biosynthesis inhibitors kynurenine (He *et al*., 2011) and yucasin (Nishimura *et al*., 2014) gradually blocked the growth-promoting temperature effect. However, as inhibition of IAA biosynthesis affected root growth already at 20°C control conditions, we regard this only as indirect evidence because it is obviously not strictly conditional. Direct evidence for an association of auxin with root thermomorphogenesis is provided by measurements of free IAA levels in root tips of five days-old seedlings grown at 20°C or 28°C. Here, higher temperature increased auxin levels in the root tip (Fig. 5B). Increased auxin levels often induce transcriptional auxin responses that can be monitored for example by imaging of auxin responsive DR5v2::tdTomato reporter activity in root tips. In confirmation of previously published data (Hanzawa *et al*., 2013; Wang *et al*., 2016; Feraru *et al*., 2019; Gaillochet *et al*., 2020), we detected significantly increased reporter activity at 28°C (Fig. 5C).

**Figure 5.**
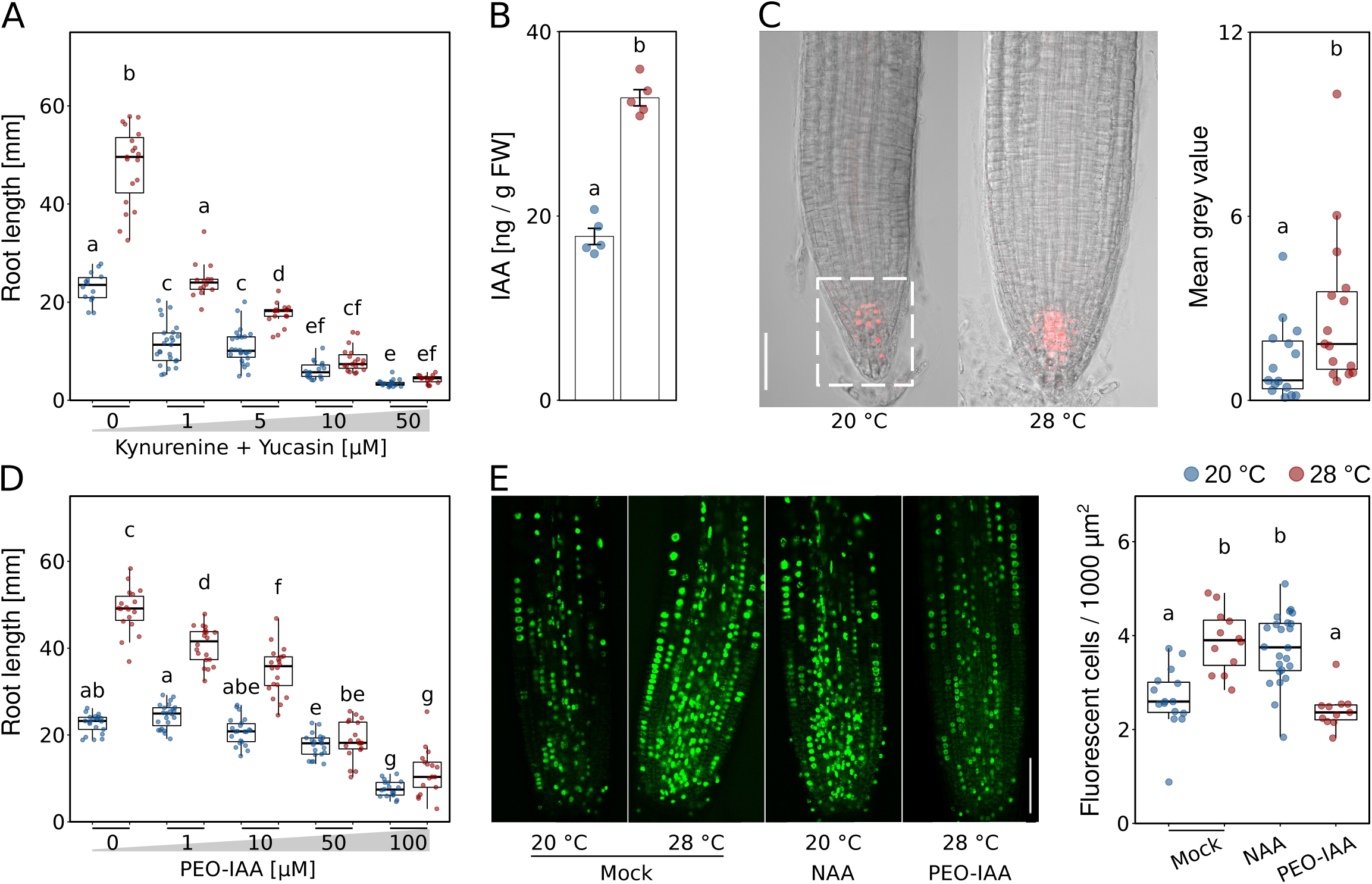
Auxin transfers temperature information to the cell cycle. **A** Root growth assay of seedlings grown for 7 days in the presence of increasing concentrations of auxin biosynthesis inhibitors kynurenine and yuccasin (n = 14-26) at the indicated temperatures. **B** Free IAA levels of root tip samples of 5 days-old Col-0 (n = 5). **C** Imaging and quantification of DR5v2::tdTomato reporter activity in root tips of 5 days-old seedlings (n = 15), scale bar = 50 μM. **D** Root growth assay of seedlings grown for 7 days in the presence of increasing concentrations of the auxin antagonist PEO-IAA (n = 17-23) at the indicated temperatures. **E** (Co-)incubation of 5 days-old seedlings for 1h with 10 μM EdU only or in combination with either 100 nM NAA or 50 μM PEO-IAA (n = 11-24), scale bar = 50 μm. **A-E** Boxplots show medians, interquartile ranges and min-max values (A,C-E). Barplot shows mean values and error bars show SEM (B). Individual data points superimposed as colored dots. Different letters denote statistical differences at P < 0.05 as assessed by one-way (B,C) or two-way ANOVA (A,D,E) and Tukey’s HSD *posthoc* test.

We then investigated temperature-induced root growth in the presence of increased concentrations of the auxin antagonist α-(Phenylethyl-2-one)-IAA (PEO-IAA), which competes with native free IAA for auxin co-receptor binding (Nishimura *et al*., 2009). While root elongation was not affected by PEO-IAA concentrations up to 10 µM at 20°C, increasing concentrations of the inhibitor within the same range gradually and significantly decreased root growth at 28°C (Fig. 5D), providing a strictly conditional phenotype. Above 10 µM PEO-IAA, seedlings grown in both temperatures were affected, albeit root lengths of seedlings grown at 28°C decreased more severely. These data support the essential role of auxin in temperature-induced root elongation proposed previously.

Auxin controls G1/S phase transition (Perrot-Rechenmann, 2010; del Pozo and Manzano, 2014). To assess a direct link between temperature, auxin and cell division rates, we next asked whether exogenous addition or inhibition of auxin would influence the number of meristematic cells entering the cell cycle. We therefore performed an EdU staining assay in the presence of the synthetic auxin naphthalene-1-acetic acid (NAA) or the auxin antagonist PEO-IAA. We found that the S-phase promoting effect of temperature alone (28°C, see also Fig. 4A) could be mimicked by adding NAA (100 nM) to the 20°C samples (Fig. 5E). Vice versa, the temperature-mediated increase in the number of cells entering the S-phase could be counteracted by adding PEO-IAA (50 μM) to the 28°C samples (Fig. 5E).

Taken together, these data indicate that auxin relays temperature information from a yet unknown thermosensor to the root apical meristem where it promotes cell proliferation in elevated ambient temperatures.

### Temperature regulates PIN expression patterns in the root tip

As auxin maxima in root apical meristems are usually generated and/or maintained by PIN-FORMED (PIN) modulated auxin distribution, we tested several *pin* mutants for their behavior in root elongation assays. We found that while *pin1-1* and *eir1-1* (a *pin2* allele) mutant roots were similar to wild type at 20°C control conditions, *pin1-1* did not respond at all to high temperatures, and *eir1-1* showed a significantly reduced temperature response (Fig. 6A). A mutant allele of PIN3 (*pin3-4*) behaved like wild type in both temperatures, suggesting that it is not required. In contrast, *pin4-2* mutants, which were likewise not significantly different from the wild type at 20°C, hyperelongated at 28°C (Fig. 6A). As such, PIN1 and PIN2 seem to act as positive regulators of temperature-induced root growth, while PIN4 inhibits excessive root growth at elevated temperatures. Importantly, these phenotypes were conditional, suggesting that these are genuine temperature effects. To further analyze the phenotypes, we used propidium iodide staining and confocal microscopy to count the number of meristematic cells in those mutants with defects in temperature-induced root elongation growth, namely *pin1-1, eir1-1*, and *pin4-2*. Along a longitudinal cell file, *pin1-1* showed a tendency for fewer meristematic cells, but these differences were not statistically significant (Fig. EV5A,B). Likewise, the *eir1-1* mutant did not differ from the wild type. In contrast, we observed an increased meristem cell number in *pin4-2* at 28°C compared to that of the wild type (Fig. EV5A,B), which may explain its long root phenotype at 28°C.

**Figure 6.**
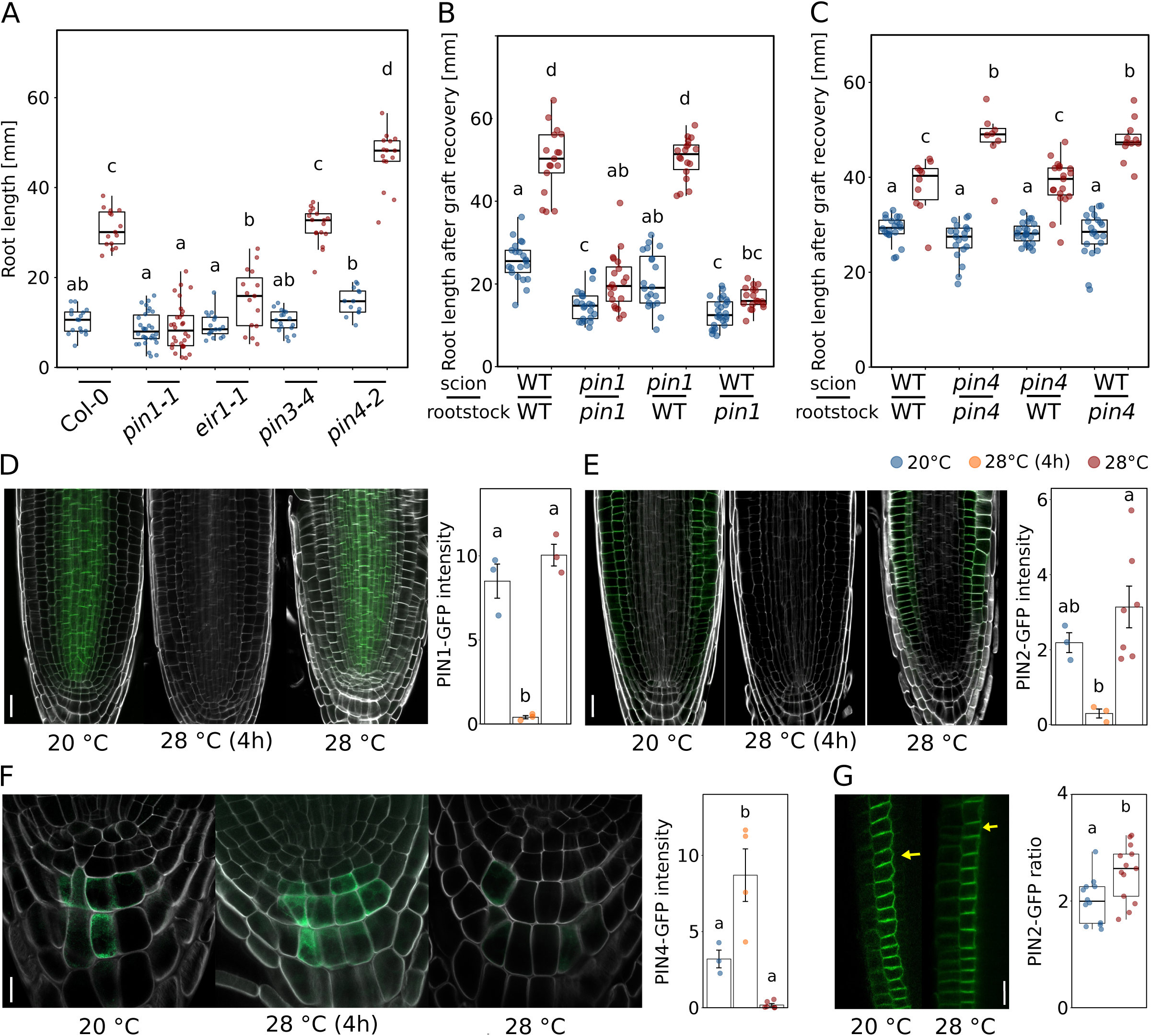
Polar auxin transport regulates root thermomorphogenesis. **A** A Root growth assay of 7 days-old seedlings (n = 13-30) at the indicated temperatures. B-C 9 days-old seedlings grown at 20°C were hypocotyl-grafted, recovered for 7 days and then cultivated at 20°C or 28°C for another 7 days (B n = 17-25, C n = 8-23). D-F Temperature effects on PIN1-GFP (D, scale bar = 20 µm), PIN2-GFP (E, scale bar = 20 µm), and PIN4-GFP (F, scale bar = 10 µm) levels. Seedlings were grown for 5 days at constant temperatures (20°C or 28°C) or grown at 20°C and shifted to 28°C for 4 h (n= 3-7), scale bar = 10 μm. G PIN2-GFP relocalization patterns (basal:lateral ratio) within cortex cells of root meristems of 5 days-old seedlings grown at the indicated temperatures. Yellow arrows mark the localizations of the lateral membrane in a cell (n = 12-13), scale bar = 20 μm. A-G Boxplots show medians, interquartile ranges and min-max values (A-C,G). Barplots shows mean values and error bars show SEM (D-F). Individual data points superimposed as colored dots. Different letters denote statistical differences at P < 0.05 as assessed by one-way (D-G) or two-way (A-C) ANOVA and Tukey’s HSD *posthoc* test.

Next, we selected one short root *pin* mutant (*pin1-1*) and one long root *pin* mutant (*pin4-2*) and asked whether these phenotypes originate in the root or may possibly be caused by long distance transport of shoot-derived auxin requiring PIN function. Micrografting showed that the observed phenotypes did only occur when *pin* mutants were used as rootstocks (Fig. 6B,C), indicating that the elevated auxin levels at high temperatures in the root tip are root-derived.

We then looked at the behavior of GFP fusion proteins of PIN1, PIN2, and PIN4 (Fig. 6A). PIN-GFP fusion proteins responded in parts strongly to temperature changes. When five days-old seedlings were shifted from 20°C to 28°C for 4 h, GFP fusion protein signals of PIN1 and PIN2 fusions in the root tip virtually disappeared (Fig. 6D,E), while PIN4 reporter levels remained unaffected (Fig. 6F). However, these PIN1 and PIN2 effects were transient, as illustrated by unchanged GFP signal intensities in seedlings constantly grown at the respective temperatures (Fig. 6D,E). It is therefore unclear whether this short-term disappearance affects root growth at all. PIN4-GFP levels in the columella showed a tendency to decrease at constant 28°C (Fig. 6F), but the differences were not statistically significant. As polar localization of PINs controls the direction of auxin flow, we also quantified ratios of basal to lateral PIN-GFP localization. For PIN2-GFP we observed an increased basal-to-lateral ratio in cortex cells of seedling roots grown at constant 28°C compared to constant 20°C (Fig. 6G). Such a shift to the basal membrane would cause auxin to be preferably transported downward into the root tip, consistent with the increase of IAA levels (Fig. 5B) as well as increased DR5v2::tdTomato reporter activity (Fig. 5C; Feraru *et al*., 2019) in the same tissues.

Collectively, the reported results indicate that elevated temperature is perceived by a yet unknown root thermosensor, which either directly or indirectly activates auxin biosynthesis and PIN-dependent polar auxin transport towards the root tip, resulting in an auxin maximum. This causes an acceleration of cell division rates in the root apical meristem, which ultimately results in increased primary root elongation at elevated temperatures.

## DISCUSSION

Research efforts in the past 15 years have generated a reasonable understanding of how shoot organs sense and respond to elevated ambient temperatures (reviewed in Quint *et al*., 2016; Delker *et al*., 2017; Balasubramanian and Casal, 2019). And although we are still lacking data from natural habitats, there is also some ecophysiological insight into the advantages of such thermomorphogenic behavior, which seems to foster evaporative cooling (Crawford *et al*., 2012; Park *et al*., 2019) and thereby photosynthetic efficiency in *A. thaliana*, but also in non-vascular plants like the liverwort *Marchantia polymorpha* (Ludwig *et al*., 2021). Understanding of thermomorphogenic root growth, however, is literally still in the dark. Just like above-ground organs, root systems are exposed to a range of different (soil) temperatures, affecting the uptake and transport of water and nutrients, root growth and development (de Lima *et al*., 2021; Koevets *et al*., 2016). Noteworthy, roots of some species are even able to actively adjust their growth in a directional manner towards or away from a temperature source in a positive or negative thermotropic manner, respectively (van Zanten *et al*., 2021).

While it is established that shoot organs can sense and respond to temperature cues, it is unclear whether the root has the same capabilities. This is, however, an important question to answer, because it affects the search for root thermosensors and signaling pathway(s). Several possibilities exist: (i) roots may be unable to sense temperature themselves and completely depend on shoot-derived long distance signaling of temperature information. In contrast, (ii) they may constitute an autonomous system that is able to sense and respond to temperature cues independent of the shoot; or (iii) they may to some degree integrate above-ground temperature information in their below-ground growth ‘decision’ process. In support of a significant influence of shoot-derived signals on root thermomorphogenesis, Gaillochet *et al*. (2020) have proposed that a shoot module consisting of HY5, which is known to coordinate above-ground with below-ground growth (Chen *et al*., 2016; van Gelderen *et al*., 2018), together with phytochromes and PIFs exerts a central function in coupling growth responses between shoot and root in a temperature context. The authors suggested that a PHY-PIF-HY5 module may activate auxin and/or gibberellins, which then act as mobile signals to regulate root growth. However, the ability of detached wild type roots to elongate more at 28°C than at 20°C (Bellstaedt *et al*., 2019; Fig. 2A, Fig. EV1B) in combination with the lack of phenotypes of either *pifQ, pif4-2*, and *hy5-51* mutant shoots grafted onto wild type rootstocks (Fig. 2B; Fig. EV2B,C) favors a scenario in which roots are to be regarded as autonomous systems that can independently sense and respond to temperature cues. This does not rule out the presence of temperature-sensitive shoot-to-root communication, possibly involving phyB (Fig. EV2D), but renders it non-essential for temperature-induced root elongation. Although intact seedlings of selected shoot thermomorphogenesis mutants did show weak root growth defects, all tested lines were able to respond to high temperature (Fig. 2C,D; Yang *et al*., 2017; Gaillochet *et al*., 2020). We therefore conclude that, although being expressed in the root, with exception of HY5 (Lee et al., 2021), the major regulators of shoot thermomorphogenesis most likely play a rather secondary role in root thermomorphogenesis, and primary regulators yet wait to be identified. In any case, this suggests the presence of a different signaling pathway.

To get a handle on such a pathway, it helps to understand which cellular process is mediating temperature-induced growth in the primary root. In the shoot, the primary cellular process promoting hypocotyl and petiole growth at elevated temperatures is cell elongation. We found that roots of seedlings grown at elevated temperatures generate substantially more cells than those grown at lower temperatures (Fig. EV3B). This temperature-sensitive increase in cell numbers seemed restricted to differentiating and mature cells, while meristematic and elongating cells were hardly affected. Since cell length appeared largely temperature-insensitive across all stereotypical root zones (Fig. 3A), the only possible explanation for exaggerated root elongation at elevated temperatures is an increase of cell division rates in the root apical meristem. Obviously, this is a fundamental mechanistic difference in the manner in which high temperature cues are translated to growth responses between shoot and root tissues, respectively.

Quantification of cells at different stages of the cell cycle supports this conclusion. We observed that at 28°C more cells entered the cell cycle (Fig. 4A), more cells transitioned from G2 to mitosis (Fig. 4B), and likewise more cells were actively dividing (Fig. 4C) in the root apical meristem in comparison to plants grown at 20°C. These data are complementing a recent elegant and detailed kinematic analysis of root growth at different temperatures, showing that between 20°C and 25°C root cell production rates increased, mainly because of an increased rate of cell division (Yang *et al*., 2017). Compared to cell elongation, cell division involves further investment into cell material, it may thus be more costly in terms of biomass. But a higher number of cells also implies more cell walls that are supporting the rigidity of the structure. While this investment may not make sense for shoot tissues that face comparably little resistance in their aerial environment, more cell walls increase physical strength (e.g., force to fracture) and therefore confer higher resistance to increased soil pressure. Obviously, this is an asset when roots explore deeper soil layers.

To understand how elevated temperatures promote primary root elongation, we need to understand how temperature information connects to the cell cycle. One such connector promised to be auxin, which (i) also acts as a messenger between thermosensing and response in shoot thermomorphogenesis (Bellstaedt *et al*., 2019), and (ii) is known to promote the entrance of cells into the cell cycle especially by activating G1-S phase genes (Perrot-Rechenmann, 2010; del Pozo and Manzano, 2014). While several groups had previously shown that auxin plays an important role in root thermomorphogenesis (Hanzawa *et al*., 2013; Wang *et al*., 2016; Feraru *et al*., 2019; Gaillochet *et al*., 2020), its mechanistic role remained rather vague. Our data now support a model where auxin connects a temperature signal with the cell cycle in the root apical meristem. At high temperatures, free IAA levels are significantly increased in the root tip (Fig. 5B,C) which contains the root apical meristem. While we do not know where and via which pathway this IAA is synthesized, our grafting data indicate that it is not coming from the shoot. Within the root, IAA may be synthesized in the root apical meristem itself or elsewhere in the root and then be transported downwards. Conditional root growth phenotypes of several *pin* loss-of-function mutants suggest the involvement of polar auxin transport in the generation and/or maintenance of this auxin maximum in the root tip, which is supported by the expression pattern of, for example, PIN2. For PIN2-GFP, we observed a quantitative shift at 28°C from the lateral to the basal membrane of cortex cells, likely resulting in auxin transport downwards into the root tip (Fig. 6G). Under the reverse fountain model (Benkova *et al*., 2003; Grieneisen *et al*., 2007), auxin is redistributed in the columella to the outer cell layers where it is transported upwards again, enabling a constant auxin flow through the root apical meristem. Redistribution in the columella requires PIN4, which also functions in generating a local sink and thereby an auxin gradient in the root tip (Friml *et al*., 2002). We observed a tendency of decreased PIN4-GFP levels in the columella at 28°C (not significant; Fig. 6F), possibly interrupting this auxin flow and instead trapping auxin in the root tip, resulting in the maximum we detected in root tips of plants grown at high temperature (Fig. 5B,C). However, *pin4-2* mutants displayed hyperelongated roots at elevated temperature (Fig. 6A), making PIN4 a negative regulator of this response, suggesting that PIN4’s role in generating an auxin maximum in the root tip is likely secondary to its role in restricting meristem size (Fig. EV5). Although its localization patterns seemed unresponsive to temperature and PIN1-GFP levels were only transiently affected by high temperature (Fig. 6D), PIN1 must be essential for this process, as its loss in *pin1-1* mutants completely abrogated temperature-induced root elongation (Fig. 6A). As a result of elevated IAA levels in the meristematic region, cell division rates are increased. This process is likely reversible as shown by the effects of exogenously added auxin or auxin antagonist on the number of S-phase cells in the root apical meristem (Fig. 5E).

### Conclusions

We propose a model in which roots can autonomously sense and respond to elevated ambient temperatures (Fig. 7). In elevated temperatures, one or several yet unknown thermosensor(s) activate(s) auxin biosynthesis in the root, most likely within or close to the root apical meristem. The generated auxin maximum in the root tip is maintained in a PIN-dependent manner and promotes the entry of cells into the cell cycle, causing an increase of cell division rates. As such, auxin functions as a gas pedal. However, the role of PIN4 seems more complex than suggested in this simple model. We find that a likely reason for the hypersensitive root elongation response of *pin4-2* mutants in high temperature is an increase in meristem size (Fig. EV5). As such, PIN4 is not only redistributing auxin in the columella, it also restricts meristem size at high temperatures. To use the same analogy as above, functional PIN4 might act as a brake pedal, restricting meristem size and thereby preventing hyperelongation of the root by extending its meristematic zone.

**Figure 7.**
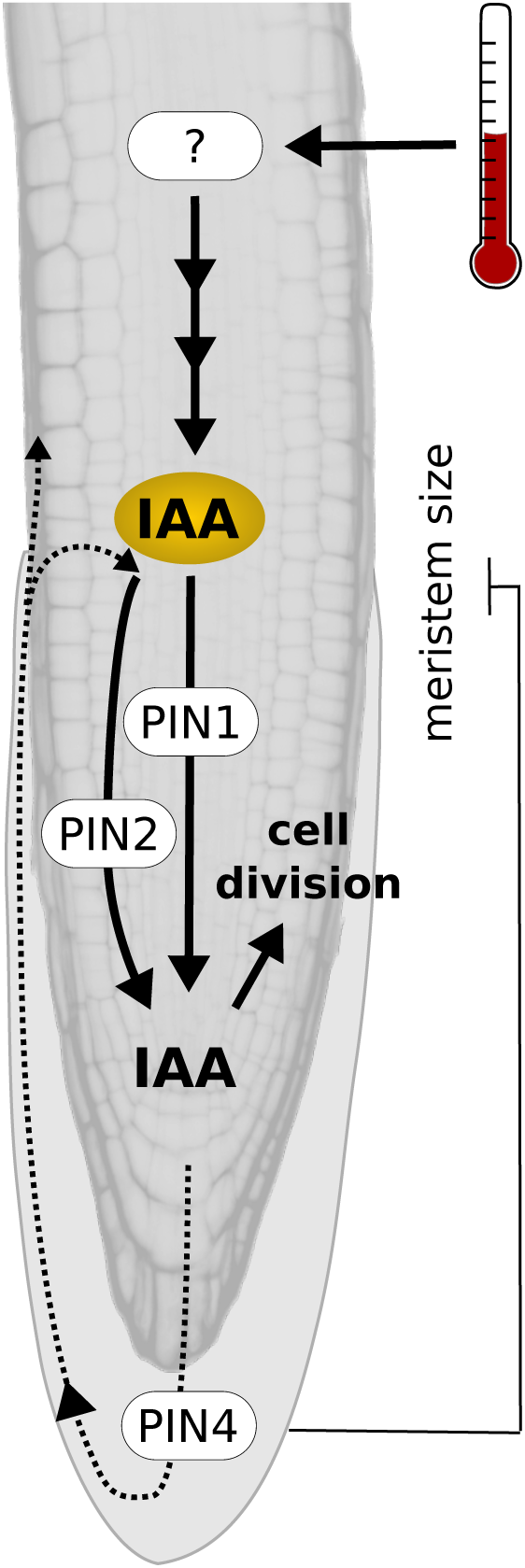
Schematic model of the major regulatory processes in the root apical meristem during root thermomorphogenesis. Elevated temperature sensed by a yet unknown thermosensor induces auxin biosynthesis, resulting in elevated auxin levels in the root tip. Polar auxin transporters PIN1, PIN2, and PIN4 help to maintain an auxin maximum in the root apical meristem which accelerates cell division rates, driving primary root elongation. In addition, functional PIN4 restricts meristem size at elevated temperature, possibly preventing excessive root growth.

Furthermore, we do not know anything about the nature of thermosensors in the root. In case thermosensing occurs in growing tissues of the roots, it would ensure short distances from the location of thermosensing to the growth-promoting cellular process, cell division, in the root apical meristem. Such a scenario would allow the primary root to determine its thermomorphogenic behavior primarily on basis of the deepest soil layers it has already penetrated. While this seems rather one-dimensional, it may be a highly versatile system. Provided that the same mechanism is active in lateral roots, it would allow primary root and lateral roots to respond to temperature independently of each other. Root plasticity in different soil temperatures may then be based on semi-independent root modules or phytomers. Alternatively, multiple thermosensors may be acting in different regions of the root, which would enable (but also require) to integrate more complex temperature information across various soil zones. If so, the root would be in need of a long distance signal transporting temperature information from various regions in the root to the root apical meristem. Auxin would be a natural candidate for this type of mobile signal.

More detailed analyses are needed to investigate via which biosynthesis pathway auxin is produced, which cell cycle components are regulated by auxin, and how other hormones cross-feed information into this network. In any case, to understand root thermomorphogenesis on a mechanistic level, we might have to refrain from trying to find parallels to shoot thermo-signaling pathways, which ultimately modulate cell elongation. Instead, it seems important to understand how temperature information influences the cell cycle in the root apical meristem.

## MATERIALS AND METHODS

### Plant material and growth conditions

Seeds of *A. thaliana* and all other species were surface sterilized, rinsed with sterile water, and then imbibed and stratified for 3 days at 4°C in sterile water before sowing. *A. thaliana* genotypes used in this study have been described previously or were obtained from the Nottingham Arabidopsis Stock Centre (NASC; http://arabidopsis.info): *cop1-6* (McNellis *et al*., 1994), *elf3-1* (N3787; Hicks *et al*., 1996), *hy5-51* (N596651; Alonso *et al*., 2003), *pif4-2* (N66043, Leivar *et al*., 2008), *pifQ* (N66049, Leivar *et al*., 2008), *35S::PIF4-HA* (Nozue *et al*., 2007), *phyABCDE* (Hu *et al*., 2013), *YHB* (P35S:AtPHYBY276H in phyA-201 phyB-5; Su and Lagarias, 2007), *phyB-1* (Reed *et al*., 1994), *cytrap* (Yin *et al*., 2014), *pin1-1* (Okada *et al*., 1991), *eir1-1* (Luschnig *et al*., 1998), *pin3-4* (Friml *et al*., 2003), *pin4-2* (Friml *et al*., 2002), PIN1::PIN1-GFP (Benková *et al*., 2003), PIN2::PIN2-GFP (Luschnig *et al*., 1998; Muller *et al*., 1998), PIN3::PIN3-GFP (Žádníková *et al*., 2010), PIN4::PIN4-GFP (Blilou *et al*., 2005). Wild type strains were Col-0 (N19992), and Ler-0 (NW20). Seeds from other species are designated as follows: *Solanum lycopersicum* (cv West Virginia 106) and *Brassica oleracea* (cv collard, NASC ID N29002). Unless stated otherwise, seedlings were grown on solid *Arabidopsis thaliana* solution (ATS, Lincoln *et al*., 1990) nutrient medium including 1 % (w/v) sucrose on vertically oriented plates under long-day conditions (16 h of light/8 h of dark) with 90 µmol m^−2^ s^−1^ photosynthetically active radiation (PAR) from white fluorescent lamps (T5 4000K).

### Root growth assays

Seedlings were placed on ATS medium, grown at 20°C or 28°C and root length was determined seven days after sowing (unless stated otherwise). Where indicated, plates were supplemented with various concentrations of auxin inhibitors kynurenine (He *et al*., 2011) and yucasin (Nishimura *et al*., 2014) or PEO-IAA (Hayashi *et al*., 2012). All measurements were based on digital photographs of plates using RootDetection (www.labutils.de) and depict the total length of the root.

For detached root assays, four days-old Arabidopsis Col-0 and *B. oleracea* seedlings, as well as five days-old *S. lycopersicum* seedlings grown at 20°C were dissected at the root-shoot junction to obtain roots only. The detached roots were then grown on vertically oriented ATS plates at either 20°C or 28°C for another four days.

### Infrared imaging of root growth

Vertically oriented ATS plates (with two days-old seedlings grown at 20°C or 28°C) were put perpendicular to the camera (Panasonic G5 with hotmirror filter replaced by an IR filter, enabling only IR light to reach the sensor; www.irrecams.de). To monitor root growth dynamics, pictures were automatically taken every hour. Derived root lengths were used to calculate root growth rates.

### Hypocotyl micrografting

Grafting was performed on seven days-old seedlings and carried out according to previously published protocols (Melnyk, 2017) and as described in Serivichyaswat *et al*. (2022). In brief, seeds were sown on ATS medium at 4°C darkness, stratified for two days at 4°C and then shifted to 20°C in a growth cabinet for another seven days under long-day photoperiods (16 h of light/8 h of dark) with 90 µmol m^−2^ s^−1^ white light (T5 4000K). Next, seedlings were grafted and recovered for seven days on a water mounted filter paper/membrane. Successfully recovered grafted plants were selected, transferred to new ATS medium and cultivated at 20°C or 28°C, respectively, under the same conditions described above for another seven days. Root growth differences were then determined by measuring the root growth between day 16 and day 23. The micrografting procedure is detailed in Fig. S2A.

### Measurement of cell length and cell number

Root cell measurements were conducted by staining seedlings with Calcofluor White (Merck, 18909-100ML-F). Briefly, seedlings were fixed in pure ethanol for 2 h to overnight, washed twice with 1x PBS, followed by a permeabilization step using 3 % Triton X-100 + 10 % DMSO in 1x PBS for 30 min to 1 h. Next, 0.1 % Calcofluor White in 1x PBS was freshly prepared and the seedlings were stained for 30 min. Subsequently, seedlings were washed twice in 1x PBS with gentle shaking. To image Calcofluor White, we used 405 nm excitation and detected signals at 425 - 475 nm. All measurements were performed on all individual cells of a consecutive cortex cell file using the ZEN 3.1 software (Zeiss) for 8 - 12 independent seedlings per experiment. The meristematic zone was defined as the zone between the quiescent centre and the last cell that did not yet double its size in comparison with the previous cell. The elongation zone followed the meristematic zone and was defined as the zone from first cell with double the size of the previous cell to the last cell before root hair bulges became visible. The following differentiation and maturation zone was defined as the zone from first cell below the first trichoblast bulge to the root-shoot junction.

### EdU staining

5-Ethynyl-2’-deoxyuridine (EdU) staining was performed with the EdU Click-488 Imaging Kit (Carl-Roth) according to the manufacturer’s instruction. Briefly, five days-old Col-0 seedlings (at ZT1, 1 h after lights on) were immersed for 1 h in liquid ATS medium containing 10 μM EdU, and fixed in 4 % (w/v) paraformaldehyde and 0.5 % Triton X-100 for 20 min. After washing twice with 1x PBS, samples were incubated in the reaction cocktail for 30 min in the dark. The reaction cocktail was then removed, and samples were washed with 1x PBS, followed by confocal microscopy with a Zeiss LSM 780 AxioObserver (excitation wavelength: 488 nm; emission wavelength: 491-585 nm). The region of interest (root meristem) was determined with the same fixed area in all measurements, and positively stained cells were counted in this area to calculate cells per 1000 μm^2^.

When stated, seedlings were co-incubated with the auxin antagonist PEO-IAA or the synthetic auxin NAA (Duchefa). Here, seedlings were stained with EdU (10 μM) + PEO-IAA (50 μM) or EdU (10 μM) + NAA (100 nM) in liquid ATS for 2-3 h (DSMO as mock) prior to fixation as described above.

### DAPI staining

At ZT2 - ZT3 (2-3 hrs after lights on), five days-old Col-0 seedlings were fixed with pure ethanol for 2 h, rinsed twice with 1x PBS, followed by a permeabilization step using 3 % Triton X-100 + 10 % DMSO in 1x PBS for 30 min. Seedlings were subsequently washed three times with 1x PBS, and then stained with 100 μg mL^−1^ 4’,6-diamidino-2-phenylindole (DAPI) in the dark for 15 min at room temperature, and processed under a Zeiss LSM 780 AxioObserver (excitation wavelength: 405 nm; emission wavelength: 425-508 nm). The region of interest (root meristem) was determined, and all stained nuclei in this area were counted by using the ImageJ software. The mitosis ratio was determined by counting cells in mitosis (condensed chromosomes visible) divided by all nuclei in this area.

### Auxin reporter assay

Five days-old Col-0 seedlings carrying the DR5v2::tdTomato reporter were fixed directly with 4 % (w/v) paraformaldehyde at room temperature, washed with 1x PBS, and kept in the dark until imaging (excitation wavelength: 561 nm; emission wavelength: 571-615 nm). Columella cells including the quiescent center were determined as the fixed area through all measurements. Mean grey values were measured by using ImageJ.

### IAA analytics

Col-0 seedlings were grown for five days as described above at 20°C or 28°C. On day 5, root tips were harvested and immediately homogenized in liquid nitrogen. Extraction of indole 3-acetic acid (IAA) was done using 50 mg homogenized material and three rounds of extraction with 400 µL, 200 µL or 100 µL of 80 % methanol, which was acidified to pH 2.4 with hydrochloric acid. In order to enhance cell rupture and extraction one steel bead of 3 mm, three steel beads of 1 mm diameter, and glass beads of 0.75 to 1 mm diameter (Carl Roth GmbH) were added to each sample and bead milling was performed for 3 × 1 min in a homogenizer (FastPrep24, MP Biomedicals). The combined extracts were centrifuged and stored on ice until measurement on the same day.

IAA was separated on a Nucleoshell RP Biphenyl column (100 mm × 2 mm, 2.1 µm, Macherey und Nagel, Düren, Germany) with the following gradient: 0-2 min: 95 % A, 5 % B, 13 min: 5 % A, 95 % B, 13-15 min: 5 % A, 95 % B, 15-18 min: 95 % A, 5 % B. The column temperature was 40°C, solvent A was 0.3 mM ammonium formate, acidified with formic acid to pH 3.0, solvent B was acetonitrile. The autosampler temperature was maintained at 4°C. Per sample 600 µL of plant extract was injected to a divinylbenzene stationary phase micro-SPE cartridge (SparkHolland B.V., Emmen, The Netherlands) at a rate of 200 µL min^-1^ where IAA is trapped by simultaneous addition of excess water (3800 µL min^-1^). Transfer from the SPE cartridge to the UPLC column was accomplished by 120 µL 20 % acetonitrile under continuous dilution with water, which gave with a final share of 2.5% acetinitrile on-column. The entire procedure was conducted on a prototype device consisting of a CTC Combi-PAL autosampler equipped with a 1 mL injection loop, an ACE 96-well plate SPE unit, a high pressure dispenser, a SPH1299 UPLC gradient pump, an EPH30 UPLC dilution pump and a Mistral column oven (all AxelSemrau GmbH, Sprockhövel, Germany). Mass-spectrometric detection of IAA on a QTrap 6500 (Sciex) was accomplished by electrospray ionization in positive mode and multiple reaction monitoring (MRM). IAA quantification was made based on transition 176/130 and was confirmed by transition 176/103 using these parameters: declustering potential: 81 V, collision energy: 27 and 46 V, cell exit potential: 8 and 11 V, respectively. For this, the ion source was heated to 450°C. Curtain gas 35 psi, ion source GS1 was set to 60 psi, GS2: 70 psi, the electrospray ion spray voltage was 5500V. IAA quantification was performed based on authentic IAA (Olchemin, Olomouc, Czech Republic).

### Confocal microscopy of PIN-GFP reporters and cytrap lines

Plants were stratified at 4°C for two days, grown on standard ½ MS media containing 1 % sucrose in a 16/8 hrs light/dark cycle at 20°C or 28°C for five days. For short term 20°C to 28°C shift experiments, plants grown at 20°C were incubated for an additional 4 h at 28°C. All plants were fixed and cleared using a previously established Clearsee-based protocol (Kurihara *et al*., 2015) modified to include Calcofluor White for staining of cell walls (Ursache et al., 2018). Briefly, plants were fixed in 4 % para-formaldehyde in 1x PBS for 1 h followed by three brief washes in 1x PBS and incubated over night in Clearsee solution containing Calcofluor White. Images were acquired on a Zeiss LSM-980 confocal system equipped with an Airyscan 2 detector using either a 40x (1.0 NA) water immersion objective or a 63x (1.4 NA) oil immersion for Airyscan images. Calcofluor White signal was detected using a 405 nm laser for excitation and an emission window from 420 - 430 nm. For GFP, excitation was achieved using a 488 nm laser and the emission window was 500 - 525nm. For RFP, excitation was achieved at 561 nm and emission collected at 580 - 620 nm. All images were acquired as non-saturated 16-bit sequential scans for further quantification.. The basal-to-lateral GFP signal of single cells in the cortical cell file was determined where indicated. GFP signal intensity was measured by using ImageJ software, and at least ten meristem cortical cells per seedling were taken into the measurement.

## ACKNOWLEDGEMENTS

The authors are thankful for the support of this work by the Deutsche Forschungsgemeinschaft (Qu 141/3-2 to MQ) as well as stipends from the Chinese Scholarship Council to HA and the Rosa-Luxemburg-Stiftung to JB. TGA thanks the Sofia Kovalevskaia program of the Alexander von Humboldt foundation and the Max Planck society for funding. GUB, AT, and BH received core funding from the Leibniz Institute of Plant Biochemistry. We would furthermore like to thank Elisabeth Beate Truernit for supporting KSB, Elena Feraru and Jürgen Kleine-Vehn for discussing unpublished data, and Jiri Friml and Dorata Jaworska for generously and repeatedly sharing seed stocks.

## AUTHOR CONTRIBUTIONS

HA, JB, KB, SB, GUB, and TGA performed the experiments. AT provided essential infrastructure, BH provided support for confocal microscopy, LEL, CD, and MQ supervised the experiments, CD generated the figures, HA and MQ wrote the manuscript. All authors edited the manuscript.

## EXPANDED VIEW FIGURE LEGENDS

**Figure EV1. Additional details on growth of Arabidopsis hypocotyl growth and elongation of dissected roots in other species grown at 20°C or 28°C**.

**A** Hypocotyl growth rates of seedling between days 2-7 were assessed every 2h by infra-red real-time imaging. Mean growth rates are shown as step-wise lines (n = 7) that were fitted with a ‘loess’ smoothing function shown as solid lines and the respective 95 % confidence intervals as grey ribbons. **B** Elongation responses of detached roots. Shoots were removed from 4 days-old *Brassica oleracea* and 5 days-old *Solanum lycopersicum* seedlings grown at 20°C and detached roots were grown for additional 4 days at 20°C or 28°C (n = 10-12). Boxplots show medians, interquartile ranges and min-max values with individual data points superimposed as coloured dots. Different letters denote statistical differences at P < 0.05 as assessed by two-way ANOVA and Tukey’s HSD *posthoc* test.

**Figure EV2: Additional hypocotyl micrografting assays and micrografting experimental setup**.

A Schematic representation of the grafting experiments shown in B-D and Figure 2B. Temperature-induced root elongation of B pif4-2 (n = 15-25), C hy5-51 (n = 27-37), and D phyB-9 (n = 15-28) mutants grafted with their corresponding WT Col-0. B-D Boxplots show medians, interquartile ranges and min-max values with individual data points superimposed as colored dots. Different letters denote statistical differences at P < 0.05 as assessed by two-way ANOVA and Tukey’s HSD posthoc test.

**Figure EV3. Changes in root and shoot cell numbers from mature embryos to 7 days-old seedlings**.

Number of cells in a consecutive cell file of **A** hypocotyls and **B** roots of mature embryos prior to germination (0 days) and in 7 days-old seedlings grown at 20°C or 28 °C. Boxplots show medians, interquartile ranges and min-max values with individual data points superimposed as colored dots (n = 5-23). Different letters denote statistical differences at P < 0.05 as assessed by one-way ANOVA and Tukey’s HSD *posthoc* test.

**Figure EV4. Root zone sizes and differentiation zone specifics**

Total length of root zones between day 2 and 7 after sowing of Arabidopsis seedlings grown in LD at 20°C or 28°C. **A** Meristem, **B** Elongation zone, **C** Differentiation zone. **A-C** Solid lines and points show mean root zone length with half-transparent ribbons denoting SEM (n = 5-7 individual roots) **D** Total number of cells in differentiation zone. Barplots show mean values, error bars indicate SEM. Individual data points are plotted as colored dots (n = 5-7). **E** Mean lengths of all cells in a consecutive cell file in the differentiation zone measured in 5-7 individual roots. Number of cells ranges from n = 5 (2 days, 20°C) to 272 (7 days, 28°C). Boxplots show medians, interquartile ranges and min-max values with individual data points superimposed as colored dots. **D**,**E** Different letters denote statistical differences at P < 0.05 as assessed by two-way ANOVA and Tukey’s HSD *posthoc* test.

**Figure EV5. Meristem sizes of *pin* mutants**

**A** Microscopic photographs of root tips of 5 days-old seedlings grown in LD at 20°C or 28°C. Yellow arrows mark the end of the meristem. Scale bar = 50 μm **B** Quantification of meristem cell numbers in consecutive cortex cell files. Boxplots show medians, interquartile ranges and min-max values with individual data points superimposed as colored dots. Different letters denote statistical differences at P < 0.05 as assessed by two-way ANOVA and Tukey’s HSD *posthoc* test.

