## Supplemental Figures for "Auxin-dependent acceleration of cell division rates regulates root growth at elevated temperature"

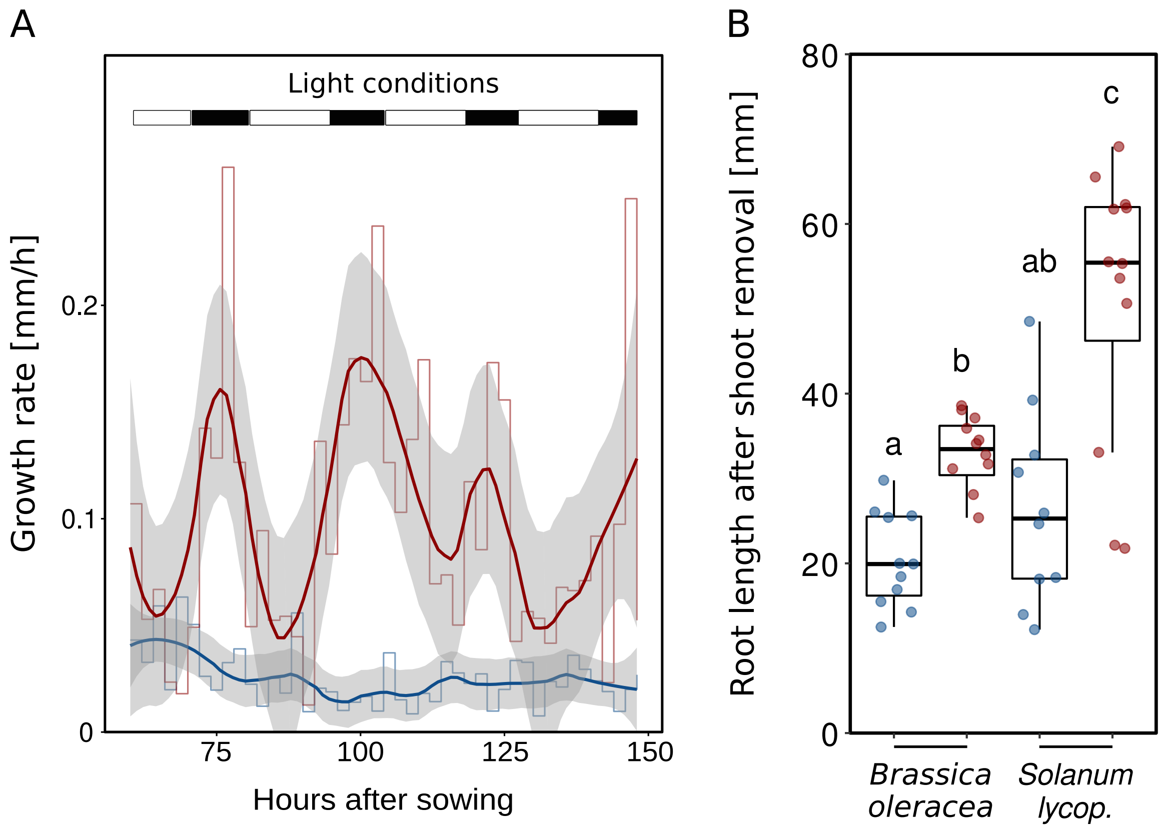


**Figure EV1. Additional details on growth of Arabidopsis hypocotyl growth and elongation of dissected roots in other species grown at 20°C or 28°C.**

**A** Hypocotyl growth rates of seedling between days 2-7 were assessed every 2h by infra-red real-time imaging. Mean growth rates are shown as step-wise lines (n = 7) that were fitted with a 'loess' smoothing function shown as solid lines and the respective 95 % confidence intervals as grey ribbons. **B** Elongation responses of detached roots. Shoots were removed from 4 days-old *Brassica oleracea* and 5 days-old *Solanum lycopersicum* seedlings grown at 20°C and detached roots were grown for additional 4 days at 20°C or 28°C (n = 10-12). Boxplots show medians, interquartile ranges and min-max values with individual data points superimposed as coloured dots. Different letters denote statistical differences at P < 0.05 as assessed by two-way ANOVA and Tukey's HSD *posthoc* test.

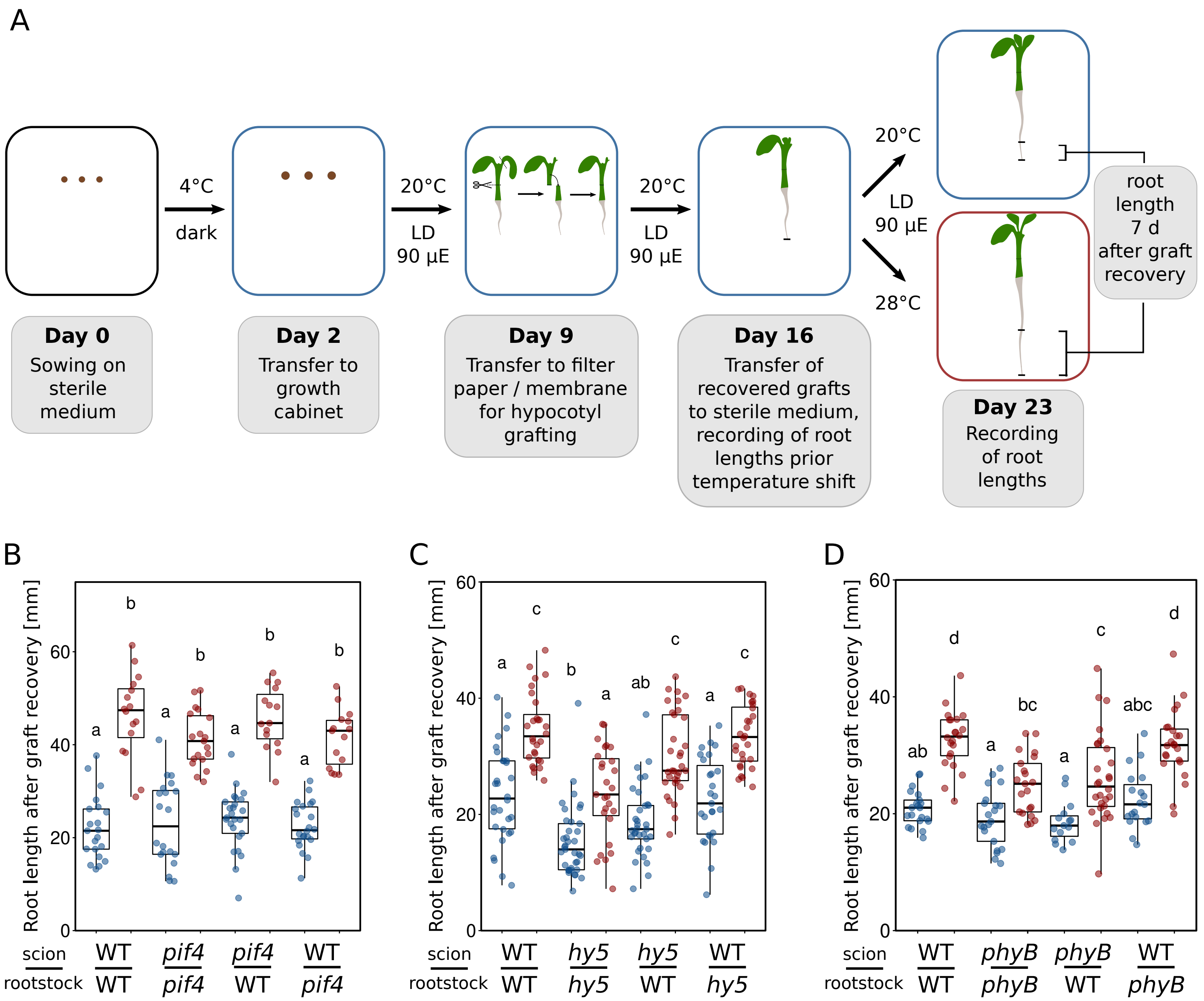


**Figure EV2. Additional hypocotyl micrografting assays and micrografting experimental setup.**
**A** Schematic representation of the grafting experiments shown in B-D and Figure 2B**.** Temperature-induced root elongation of **B** *pif4-2* (n = 15-25), **C** *hy5-51* (n = 27-37),and **D** *phyB-9* (n = 15-28) mutants grafted with their corresponding WT Col-0. **B-D** Boxplots show medians, interquartile ranges and min-max values with individual data points superimposed as coloured dots. Different letters denote statistical differences at P < 0.05 as assessed by two-way ANOVA and Tukey's HSD *posthoc* test.

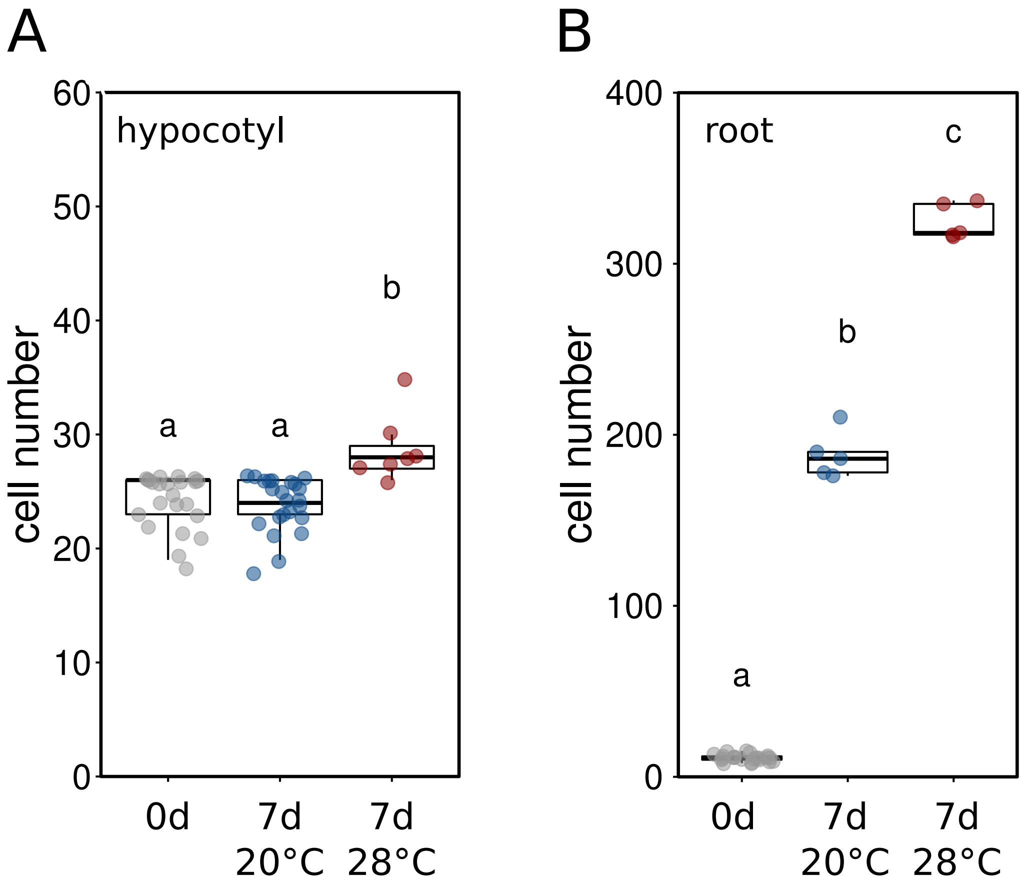


**Figure EV3. Changes in root and shoot cell numbers from mature embryos to 7 days-old seedlings.**

Number of cells in a consecutive cell file of **A** hypocotyls and **B** roots of mature embryos prior to germination (0d) and in 7 days-old seedlings grown at 20 or 28 °C. Boxplots show medians, interquartile ranges and min-max values with individual data points superimposed as coloured dots (n = 5-23). Different letters denote statistical differences at $P < 0.05$ as assessed by one-way ANOVA and Tukey's HSD *posthoc* test.

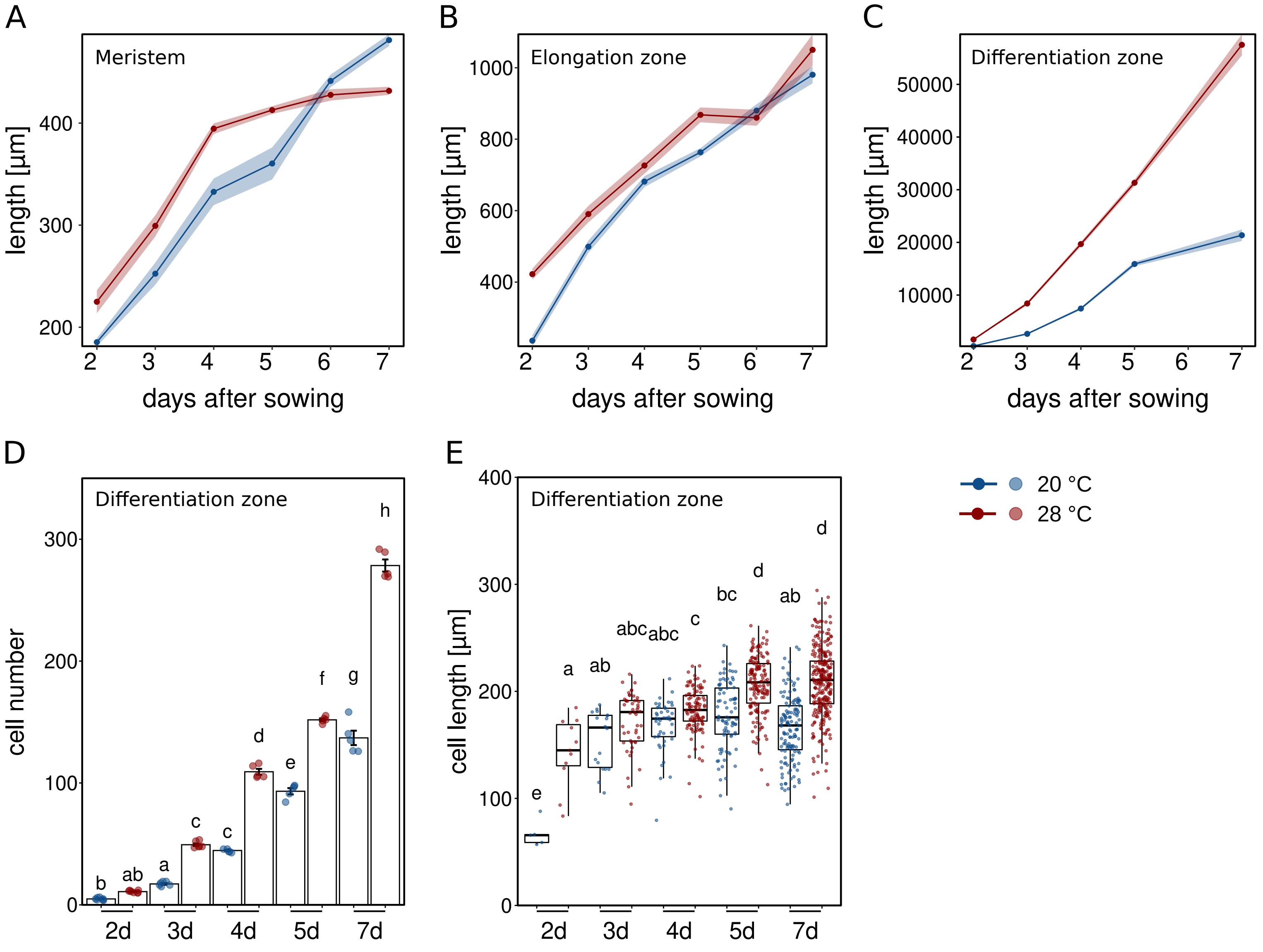


**Figure EV4. Root zone sizes and differentiation zone specifics.**
Total length of root zones between day 2 and 7 after sowing of Arabidopsis seedlings grown in LD at 20°C or 28°C. **A** Meristem, **B** Elongation zone, **C** Differentiation zone. **A-C** Solid lines and points show mean root zone lengths with half-transparent ribbons denoting SEM (n = 5-7 individual roots). **D** Total number of cells in differentiation zone. Barplots show mean values, error bars indicate SEM. Individual data points are plotted as colored dots (n= 5-7). **E** Mean lengths of all cells in a consequtive cell file in differentiation zone measured in 5-7 individual roots. Number of cells ranges from n = 5 (2d, 20°C) to n = 272 (7d, 28°C). Boxplots show medians, interquartile ranges and min-max values with individual data points superimposed as colored dots. **D,E** Different letters denote statistical differences at P < 0.05 as assessed by two-way ANOVA and Tukey's HSD *posthoc* test.

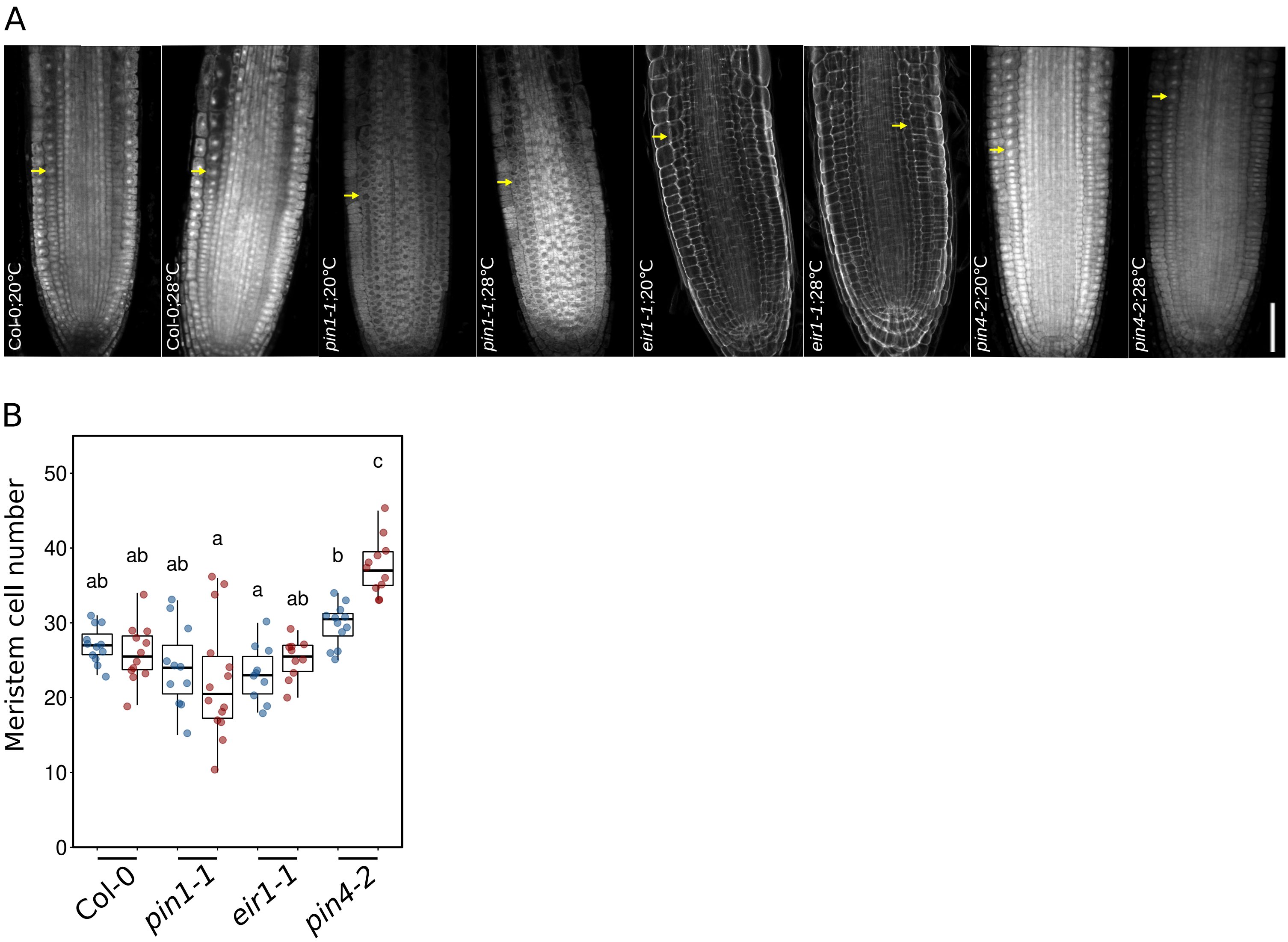


**Figure EV5. Meristem sizes of *pin* mutants.**

**A** Microscopic photographs of root tips of 5 days-old seedlings grown in LD at 20°C or 28°C. Yellow arrows mark the end of the meristem. Scale bar = 50 µm **B** Quantification of meristem cell numbers in consecutive cortex cell files. Boxplots show medians, interquartile ranges and min-max values with individual data points superimposed as colored dots. Different letters denote statistical differences at P < 0.05 as assessed by two-way ANOVA and Tukey's HSD posthoc test.
